# Intrinsic and circuit mechanisms of predictive coding in a grid cell network model

**DOI:** 10.1101/2025.04.11.648301

**Authors:** Inayath Shaikh, Collins Assisi

## Abstract

Grid cells in the medial entorhinal cortex (MEC) fire at the vertices of a hexagonal lattice, forming an allocentric code for the animal’s current position. Recent studies have identified a class of grid cells that represent locations ahead of the animal. How do these predictive representations emerge from the wetware of the MEC? We developed a detailed conductance-based model of the MEC network, constrained by empirical data on the biophysical properties of stellate cells and the topology of the MEC network. The model revealed two mechanisms by which grid cells can signal future locations. First, hyperpolarizing inhibition from interneurons activates HCN channels in stellate cells, whose slow kinetics maintain a depolarizing influence after inhibition ends, advancing spike timing and shifting the inferred position forward by ~5% of a grid field diameter. Second, introducing asymmetry into the inhibitory connectivity, by skewing the Gaussian profile of interneuron-to-stellate connections, causes inhibition to rise more steeply and enables earlier spiking, advancing the inferred position by up to ~25%. A corollary of our model is that the extent of the predictive code changes monotonically along the dorsoventral axis of the MEC, following the experimentally measured dorsoventral gradient in HCN time constants.

## 1 Introduction

Grid cells found in the medial entorhinal cortex (MEC) of mammals are spatially tuned neurons that spike at the vertices of a triangular lattice that tiles space [Hafting et al., 2005; Gardner et al., 2019]. Across different environments and even during sleep, grid cells maintain consistent phase relationships with each other and, compared to place cells, undergo minimal remapping [Fyhn et al., 2007; Gardner et al., 2019]. Grid cells are therefore thought to provide an allocentric representation of space, a coordinate system anchored to external landmarks, not to the animal itself. This coordinate system allows the animal’s trajectory to be mapped as it moves. By continuously tracking the instantaneous direction of movement and integrating the speed of travel, circuits in the brain can presumably locate the animal within a two-dimensional environment. Along the dorsoventral axis of the MEC, grid cells are organized into modules characterized by increasing grid scales (lattice constants) and larger grid field sizes. This organization is similar to a ruler with markings that repeat periodically on multiple scales, such as millimeter, centimeter, and meter markings[Stensola et al., 2012]. The population activity of grid cells can be used to infer the location of the animal with remarkable precision on a length scale that exceeds the largest lattice constant [Sreenivasan and Fiete, 2011; Fiete et al., 2008].

Grid cells represent not only present locations, but also past and potential future positions of the animal [Ouchi and Fujisawa, 2024; Chaudhuri-Vayalambrone et al., 2023]. Theoretical models of grid cell networks have proposed a process termed ‘linear look-ahead’ [Kubie and Fenton, 2012; Erdem and Hasselmo, 2012]. This process involves selective experience-dependent strengthening of connections between direction-conjunctive grid cells that share similar heading directions. As a result, the location inferred from the collective activity of these cells places the animal further ahead of its current location on a linear trajectory. Recent experiments have revealed cells called predictive grid cells in layers II and III of the MEC that also place the animal in locations further along its trajectory, rather than at its current physical location[Ouchi and Fujisawa, 2024]. More recently, a peculiar predictive representation was discovered in which the activity of the grid cells appears to periodically scan locations along the path of the animal, swinging alternately to the left and right [Vollan et al., 2025; Ji et al., 2025]. What are the mechanisms underlying predictive coding in the MEC? Can the intrinsic properties of grid cells and inhibitory interneurons contribute to the emergence of predictive coding in MEC?

To address these questions, we developed a detailed network model composed of realistic conductance-based stellate cells (a majority of grid cells in layer II of the MEC are stellate cells) and inhibitory interneurons [Pastoll et al., 2012; Wang and Buzsáki, 1996]. The hexagonal symmetric activity patterns of the grid cells are phase-shifted relative to each other and can be mapped to a low-dimensional toroidal manifold [Gardner et al., 2022]. As the animal moves, the state of the network follows a continuous trajectory on the surface of this torus. This low-dimensional attractor manifold arises from the interactions between neurons within the network. Several network models have been proposed to explain how spatiotemporal patterns constrained to a torus emerge in the MEC network [Giocomo et al., 2011a]. Among these, we focus on Continuous Attractor Network (CAN) models. These models typically use rate-based neurons or integrate and fire neurons organized spatially with distance-dependent connectivity [Burak and Fiete, 2009; Fuhs and Touretzky, 2006; Navratilova et al., 2012; Kang and DeWeese, 2019]. The connectivity kernel of neurons in this network approximates a center-surround topology, with neurons near any given grid cell receiving excitatory input (or reduced inhibition) from that cell, while neurons farther away receive inhibitory input. The connectivity is radially symmetric and results in activity patterns that peak at the vertices of a triangular lattice. Although different CAN models of grid cells share several core properties, they differ in specific details. These include whether excitatory principal neurons and inhibitory interneurons are treated as separate populations and whether the network nodes consist of spiking neurons or rate-based neurons. However, no existing model fully integrates the biophysical details of conductance-based neurons.

Using a Continuous Attractor Network (CAN) model as a framework, we incorporate the biophysical details of stellate cells and inhibitory interneurons, arranged in a network motif commonly found in the medial entorhinal cortex (MEC) [Witter et al., 2017; Neru and Assisi, 2021]. This approach allowed us to uncover the mechanistic basis of several experimentally observed phenomena. Our model network, similar to grid cells recorded *in vivo*, exhibits a depolarizing ramp as the rat enters a grid field [Domnisoru et al., 2013]. We found that this depolarization arises from gradual disinhibition mediated by inhibitory interneurons. Disinhibition, combined with the rebound properties of HCN channels, contributes to the predictive bias observed in our model. Additionally, when we simulated the knockout of HCN channels in our model, the velocity gain of the attractor was reduced, leading to an expansion of grid scales and field sizes. This outcome is consistent with experimental observations *in vivo* [Giocomo et al., 2011b].

## 2 Results

### 2.1 Biophysically detailed conductance-based grid cell attractor model

To build a continuous attractor model of grid cells, we used conductance-based neuron models of stellate cells and interneurons [Acker et al., 2003; Wang and Buzsáki, 1996]. These neuron models capture the rich intrinsic dynamics observed in MEC layer II stellate cells and fast-spiking inhibitory interneurons. Stellate cells show post-inhibitory rebound, subthreshold membrane potential oscillations, resonance, and depolarizing sag (Figure 1a). The timing of the rebound spike and the depolarizing sag is determined by the strength of a brief hyperpolarizing input (Figure 1a), top panels). Subthreshold oscillations (Figure 1a, bottom left panel) emerge before the onset of a spike (not shown). In response to a subthreshold oscillatory drive with a monotonically increasing frequency, the model stellate cell exhibited a resonance peak at a frequency close to that of the subthreshold oscillations mentioned above (Figure 1a, bottom right panel). These properties are due to the presence of a hyperpolarization-activated depolarizing current (I_*h*_) and a persistent sodium current (I_*NaP*_) in addition to leak and spiking currents (I_*L*_, I_*K*_, I_*Na*_). An interplay between I_*h*_ and I_*NaP*_ is needed to generate the membrane potential oscillations observed here [Fransén et al., 2004]. In addition, these oscillations fall squarely within the range of theta oscillations (4-12Hz) [Colgin, 2013].

**Figure 1.**
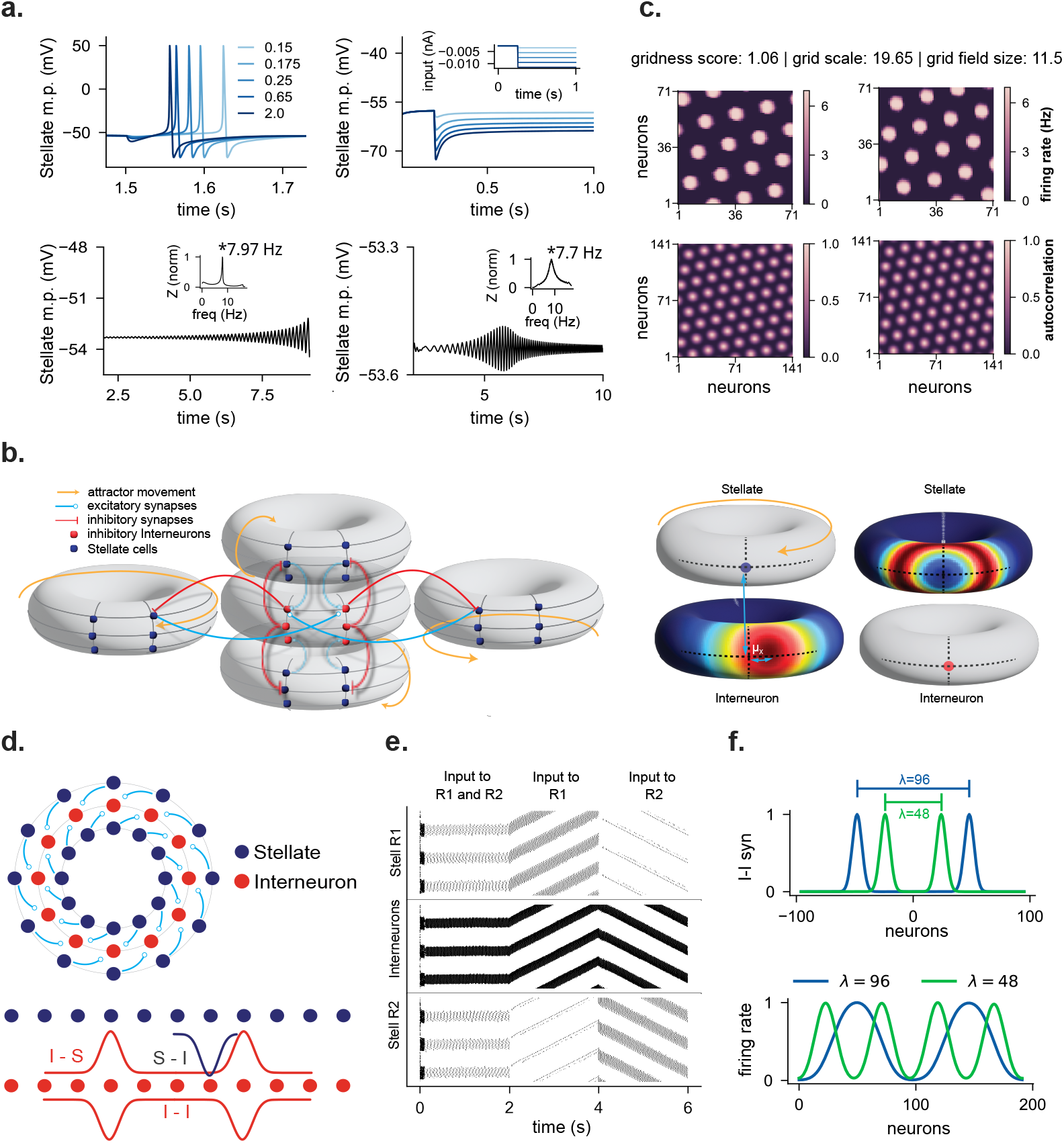
1D and 2D grid cell network and activity pattern. **(a) Stellate cell intrinsic dynamics.**In clockwise direction (1) Post-inihibitory rebound as a result of hyperpolarizing pulse input of increasing intensity. (2) Depolarizing sag in response to increasing hyperpolarizing input. (3) Membrane potential resonance in response to chirp input. inset, normalized impedance for the same trace. (4) Subthreshold oscillations in the membrane potential oscillations. inset, normalized impedance. **(b) 2D network structure** for four sheets of stellate cells and one sheet of inhibitory interneuron. The opposite edges of each sheet are connected to form a torus. The middle schematic shows the offset in the output from stellate sheet representing the east direction to interneuron sheet. The right schematic depicts center-surround connectivity from interneuron to stellate sheet. The same connectivity is used for mutual connections among interneurons (not shown). **(c) 2D network activity** of stellate cells moving in east (left) and north (right) direction, along with their autocorrelation (bottom). **(d) 1D ring network**. Two stellate rings and one interneuron ring with 192 neurons in each ring. The interneuron ring is mutually connected with a centre-surround connectivity (I-I, bottom). The same connectivity is projected onto both the stellate rings (I-S, bottom). The connection from the two stellate rings to the interneuron ring has an offset in opposite directions representing left and right movement (S-I, bottom). **(e) Raster plot for 1D network**. Top, Stellate ring 1 (R1) with clockwise offset. Middle, Interneuron ring. Bottom, Stellate ring 2 (R2) with counterclockwise offset. For the first 2 seconds, both stellate rings receive DC inputs. Then, Stellate R1 receives DC input from 2-4 seconds, followed by Stellate R2 from 4-6 seconds. **(f) Grid scale and field sizes** can be increased (bottom) by increasing the distance between two peaks of the center-surround connectivity (top)**Figure 1—video 1**. 2D Grid cell activity pattern

The network model consisted of five neuronal sheets: four sheets of stellate cells and one sheet of interneurons. We modeled each sheet with 71 × 71 neurons. Neurons at the opposite edges of each sheet were coupled resulting in a network organized as a torus. Stellate cells were not directly connected to each other, but interacted through interneuron-mediated disynaptic inhibition, as illustrated in Figure 1b [Couey et al., 2013]. Inhibitory interneurons provided radially symmetric input to other interneurons within their respective neural sheet. Similarly, the interneuron-to-stellate connectivity was also radially symmetric, with each interneuron extending connections to a ring of stellate cells (Figure 1b right). This topographic organization and radial symmetry in inhibitory connectivity resulted in hexagonal activity patterns across both the stellate and interneuron sheets. This architecture reflects the anatomical organization of layer II of the MEC, where stellate cells form spatially segregated clusters separated by patches of pyramidal neurons [Gu et al., 2018], interact primarily through inhibitory interneurons rather than through direct mutual excitation [Couey et al., 2013; Witter et al., 2017], and receive spatially organized synaptic input from parvalbumin interneurons whose axonal projections are concentrated at the scale of individual grid-cell clusters [Huang et al., 2024]. Layer II pyramidal cells, which form a largely separate microcircuit connected via distinct interneuron populations, are not included in the model, and our results and predictions are therefore specific to the stellate-cell grid-cell population.

The connectivity kernel used for the interneuron-interneuron (w^*II*^) connection weights was defined as follows,

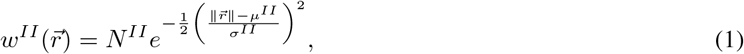

where µ^*II*^ > 0 and 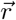 is a vector that extends from one neuron to the other on the neural sheet. The same connectivity profile projected from the interneuron to the stellate sheet (w^*IS*^). Excitatory input from the stellate cells to the interneurons was concentrated around a circular region on the interneuron sheet, with an offset in one of four cardinal directions. Activating a stellate sheet drove the grid pattern along the torus in the direction determined by this offset. The connectivity kernel for stellate-to-interneuron (w^*SI*^) interactions was expressed as:

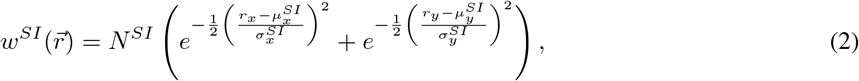

where 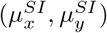determined the offset direction from the stellate to the interneurons, which influenced the movement direction of the activity bumps when that specific stellate sheet was activated.

The network topology closely resembled that described in Kang and DeWeese [2019], with key differences including mutual inhibition within the interneuron sheet, no mutual excitation between stellate cells, and edge connections forming a toroidal structure for each sheet. The radially symmetric interneuron-interneuron connections resulted in a grid-like pattern of activity (Figure 1c). By selectively stimulating the stellate cell sheets with speed-dependent depolarizing inputs, we were able to translate the grid attractor to match a simulated trajectory in 2-D space (Figure 1—video 1).

To facilitate subsequent computational analyses, we reduced the two-dimensional toroidal network to a one-dimensional ring structure. This simplified model consisted of one ring of interneurons sandwiched between two rings of stellate cells. The connectivity in this ring was derived by taking a cross-section of the torus along a principal grid axis. This can be considered equivalent to restricting the trajectory to a principal axis of the hexagonal grid. The resulting tuning curve of the grid cells exhibited spatial periodicity. The connectivity kernel for interneuron-interneuron interactions 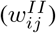, restricted to the ring, was defined based on the angular distance (Δθ_*ij*_) between neurons i and j as:

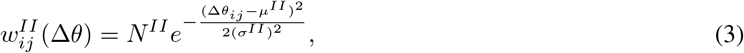

where

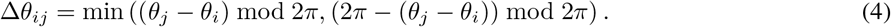

This center-surround connectivity profile was characterized by parameters µ^*II*^, which represents the distance between the peaks, and σ^*II*^, which defines the width of the peak. The same connectivity profile was applied to the interneuron-stellate connections 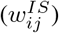, as shown in Figure 1d (bottom).

The output of stellate rings to interneurons followed a shifted Gaussian profile. The shift indicated clockwise or counterclockwise movement (+µ^*SI*^ or− µ^*SI*^, respectively) around the ring. The weights from neuron i (at angular position θ_*i*_) to neuron j (at angular position θ_*j*_) were given by:

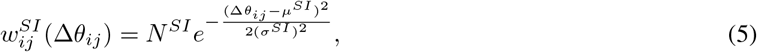

with

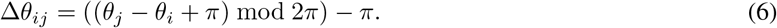

This shifted Gaussian connectivity allowed directional propagation of activity patterns within the network (Figure 1e). Grid scale and grid field sizes can be tuned by adjusting µ^*II*^ and µ^*IS*^, which determine the distance between the peaks of the centre-surround connectivity profile (Figure 1f).

### 2.2 Grid cell network can accurately path integrate 1D trajectories

Continuous attractor models of grid cells conceptualize the network as a topographically organized sheet of neurons, where adjacent areas in physical space correspond to neighboring regions in the neural network. This organization presumes a mapping between the animal’s trajectory in real space and the continuous movement of the hexagonal attractor pattern across the neural sheet. The continuity of animal movement in physical space (since animals cannot teleport across space) manifests itself as the continuous movement of the attractor pattern on the neural sheet. Some evidence supports the existence of this movement in the entorhinal cortex [Gu et al., 2018]. Attractor networks typically model the animal’s speed as a depolarizing input to grid cells, with the magnitude of this input often scaling linearly with speed. This linear relationship is consistent with observations of many speed cells, whose firing rates increase linearly with the animal’s velocity[Kropff et al., 2015]. However, the representation of speed in the brain is more complex than a simple linear model suggests. Speed cell responses can be diverse, with mostly monotonic but sometimes nonmonotonic relationships to speed, and slopes that vary widely across different speed ranges[Hinman et al., 2016]. Furthermore, the speed-tuning curves adapt to changes in the shape and size of the environment, indicating a dynamic response to the environment of the animal [Munn et al., 2020].

Accurate spatial navigation, especially in the absence of external cues, requires animals to integrate their speed and direction of motion. This process, known as path integration, relies on the activity of grid cells. A crucial property of this system is that the grid cell population activity at a specific location must be independent of the path taken to reach that point. This path invariance requires a linear relationship between the equations that govern the activity of the grid cell and the velocity of the animal [Issa and Zhang, 2012]. However, the neural mechanisms underlying this process present a challenge. Stellate cells do not respond linearly to changes in input magnitude, unlike rate-based neurons. In addition, in the network modeled here, the propagation speed of activity across the topographically organized ring network of stellate cells did not increase linearly with increasing depolarizing input.

To address this non-linearity and ensure accurate path integration, we mapped each value of speed s measured in (rad/s) to its corresponding depolarizing drive I = f(s) (orange curve, Figure 2a). The magnitude of this depolarizing drive remained within the sub-nanoampere range used for somatic current-clamp characterization of stellate cell and interneuron excitability in the studies from which our conductance parameters were derived [Alonso and Klink, 1993; Dickson et al., 2000; Fransén et al., 2004; Acker et al., 2003], indicating that the drive required for accurate path integration in our model is biologically plausible. This was done by first measuring the resulting attractor speed for a range of input currents, yielding the inverse relationship s = f ^*−*1^(I) (Figure 2a, blue curve). Since f ^*−*1^(I) increased monotonically, we could invert it to determine the magnitude of the depolarizing input that would be delivered uniformly to all stellate cells for a given speed. By applying this inverse transformation, we effectively linearize the system’s response, ensuring that bump movement becomes linear in relation to input velocities (Figure 2b). Interestingly, this computational solution finds support in observations. Experiments have revealed similar responses in speed-tuned head direction (HD) cells of the MEC in response to increased animal speed (Kropff et al. [2015], Extended Data figure 4b). This parallel between our model’s velocity processing mechanism and observed neural activity suggests that the brain may employ comparable strategies to achieve linear path integration despite the inherent nonlinearities in the responses of individual neurons and the activity of the network.

**Figure 2.**
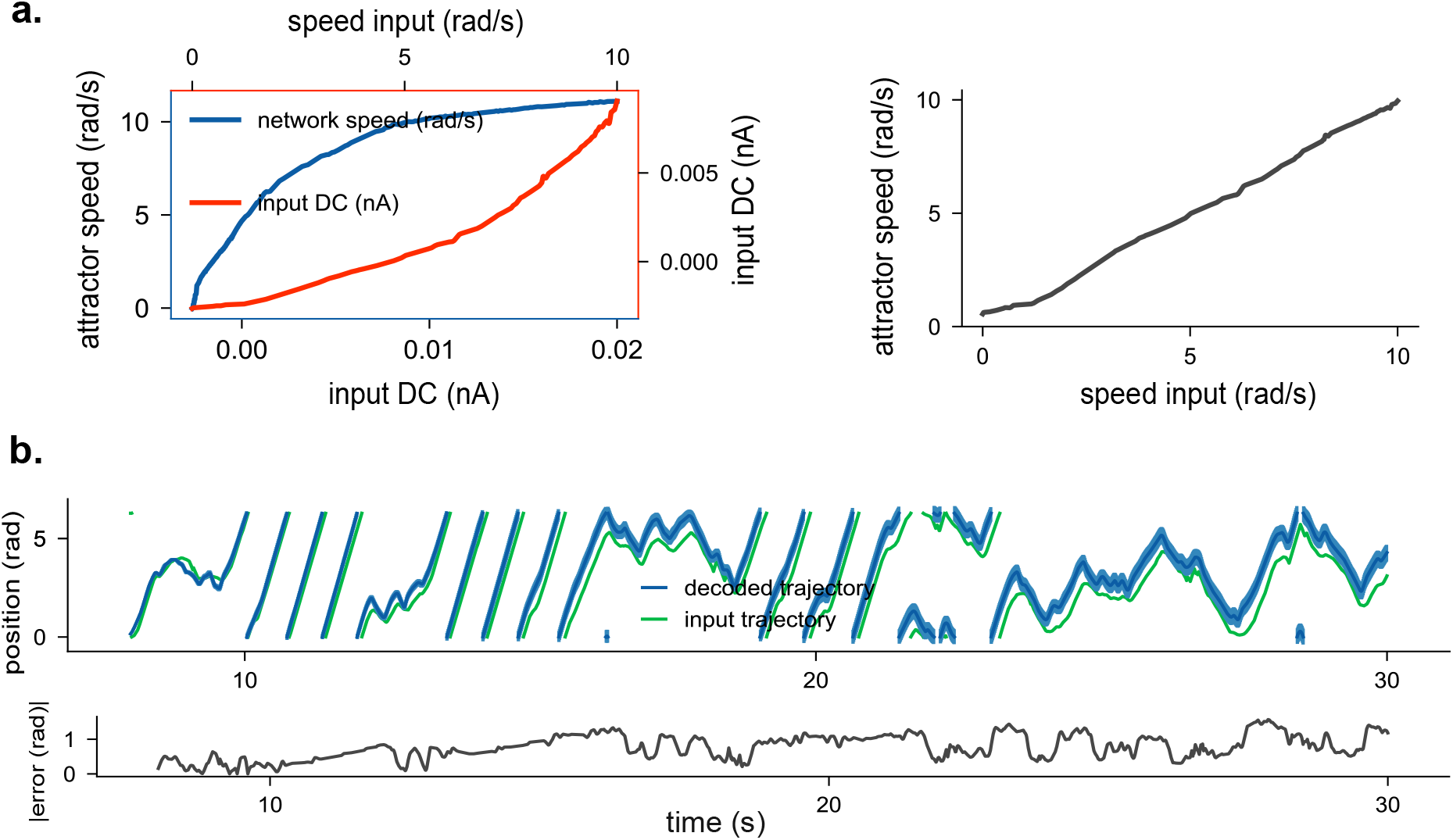
Path Integration in 1D network. **(a)** Left, speed of the network in response to increasing DC inputs to the stellate ring (in blue) and Velocity integration (V-I) curve used for path integration (in orange). Notice that the V-I curve is the inverse of the attractor speed - Input DC relation. Right, the resulting relation between attractor speed and input speed after applying the Velocity integration curve. **(b) Path integration** of a randomly generated circular trajectory (green). A population vector-based decoder is used to decode positions from the neural activity (blue). Bottom, absolute error for the decoded trajectory at different points in the simulation.

We simulated one-dimensional trajectories in a circular environment with statistical features that resembled experimentally recorded trajectories of rats [George et al., 2024]. Stellate cells in the clockwise and counterclockwise rings spiked at spatially periodic locations. The firing fields of different stellate cells were shifted with respect to each other. We used a population vector-based decoder to infer the position of the animal given the collective activity of stellate cells [Stemmler et al., 2015]. We then compared the inferred trajectory of the animal with the actual path and found that it tracks the location of the animal accurately over behaviorally relevant timescales (the simulated trajectory of the animal extended over 30 seconds).

### 2.3 Disinhibition generates depolarizing ramps

The network architecture simulated here, along with the mechanisms underlying the emergence of grid-like activity in stellate cells, closely resembles that found in attractor models [Khona and Fiete, 2022]. A key prediction of the attractor model is that grid cell firing fields result from the gradual ramping of depolarization due to synaptic inputs from the rest of the network. The grid fields are centered around the peaks of these depolarizing ramps. *In vivo* recordings confirm this prediction [Domnisoru et al., 2013]. How does the simulated network generate such ramp-like potentials? Mutual excitatory connections between stellate cells are believed to be sparse [Couey et al., 2013; Witter et al., 2017]. Thus, any depolarizing synaptic input must come from an external source that selectively excites sub-populations of stellate cells in a location-dependent manner [Neru and Assisi, 2021]. In our model, this selectivity is absent as the only external excitatory input modulates all stellate cells uniformly. However, neurons in the network still exhibit grid-like firing fields and accurately integrate the animal’s trajectory.

Following the analysis by Domnisoru et al. [2013], we removed spikes from the membrane potential of neurons in our model and applied a low-pass filter (cutoff at 3 Hz) to test if our model generated ramp-like potentials observed in experiments (see Methods and Materials). As the trajectory traversed the firing field of a stellate cell, the spike-free membrane potential gradually increased, then decreased as the trajectory left the field (Figure 3a, top). These changes in membrane potential were due to disinhibition, with inhibitory inputs to the stellate cell decreasing as the animal entered the field and increasing as it exited (Figure 3c).

**Figure 3.**
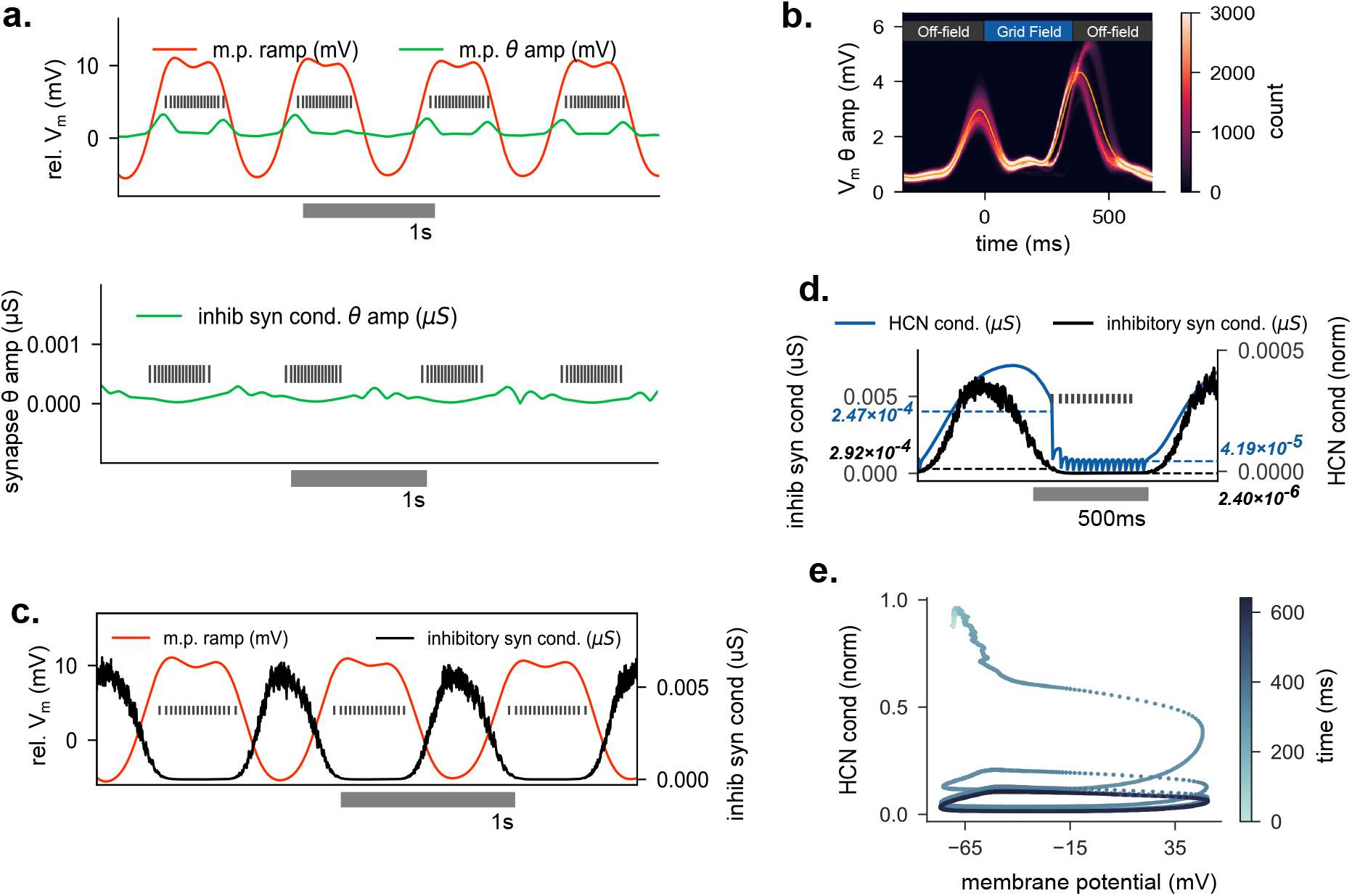
Membrane potential dynamics along grid field **(a) Membrane potential ramps and theta modulation.**Top, Membrane potential ramps (orange) compared with its theta amplitude envelope (green). Spikes demarcating the grid fields are shown in grey. Bottom, Theta amplitude envelope of the overall inhibitory synaptic conductance received by the same cell. **(b)** Time series histogram of membrane potential theta modulation of all stellate cells. The average is marked in yellow and the location of the grid field is denoted above. **(c) Disinhibition for ramp formation**. Inhibitory synaptic conductance (in black) to stellate cells shows disinhibition as a mechanism for membrane potential ramp formation (orange). **(d) First spike as postinhibitory rebound**. Inhibitory synaptic conductance (black) and HCN conductance (blue) along the grid field (gray). Annotations denote the conductance values at the first (green) and the last spike (orange) of the field. At the onset of the field, as the inhibition wanes, the HCN conductance is higher when compared with the end of the field, leading to post-inhibitory rebounds. **(e)** Phase plot for HCN conductance and membrane potential for 300ms before the onset of the field to the end of the field.

Furthermore, intracellular recordings have shown that changes in membrane potential within the firing field are not accompanied by changes in theta oscillation amplitude [Domnisoru et al., 2013]. To test this in our model, we filtered the spike-free potential in the theta band (4–12 Hz) and plotted its amplitude (see Methods and Materials, Domnisoru et al. [2013]). Consistent with experimental results, the variations in theta oscillation amplitude in our model were an order of magnitude lower than the ramp potential caused by disinhibition. Although theta power was low both within and outside of the firing field, we observed distinct low-amplitude theta modulation (Figure 3b) near the borders of grid fields. These flanking peaks, indicating entry in and exit out of a grid field, were not the result of theta-frequency synaptic input from interneurons, but were intrinsically generated by stellate cells (Figure 3a, bottom), due to the combined effects of HCN and Na_*p*_ channels [Alonso and Llinás, 1989].

As a consequence of disinhibition and the presence of HCN channels, the first few spikes of grid fields in our model were driven by HCN-driven depolarizing currents resulting in post-inhibitory rebound spikes. HCN channels, upon inhibition of stellate cells by interneurons, activate, reaching a peak conductance value 100-200 ms before the onset of the grid fields (Figure 3d). As inhibition subsides, the excitatory drive from HCN-dependent depolarization outweighs the inhibitory input from interneurons, leading to rebound spikes (Figure 3d,e).

### 2.4 HCN scales the grid cell attractor by lowering the velocity gain

HCN knockout experiments in mice show an expansion of grid scales and grid fields across the dorso-ventral axis of the MEC [Giocomo et al., 2011b]. In contrast, in our model, HCN knockout leads to a reduction in the size of the activity bumps and shows no significant change in the distance between the peaks (Figure 4b,e). How can these model results be reconciled with experimental observations? The modeled grid attractor network is local, radially symmetric, and translationally invariant. Our model, like all attractor models, assumes a distance-preserving mapping between real space and the space of neurons. In experiments, one measures the activity of a single stellate cell network. As an animal moves, the attractor moves as well. A particular neuron fires when a peak of the spatially periodic attractor passes over it. There are two potential ways to change the measured width of an activity bump: either by recruiting more neighboring neurons so that the activity bump becomes wider or by slowing the movement of the bump, effectively decreasing the velocity gain of the attractor (Figure 4a). Our model suggests that HCN knockout leads to a slowdown of the attractor along the neuronal sheet. This slowing also results in an increase in the measured distance separating two activity bumps. What are the mechanisms underlying this slowdown?

**Figure 4.**
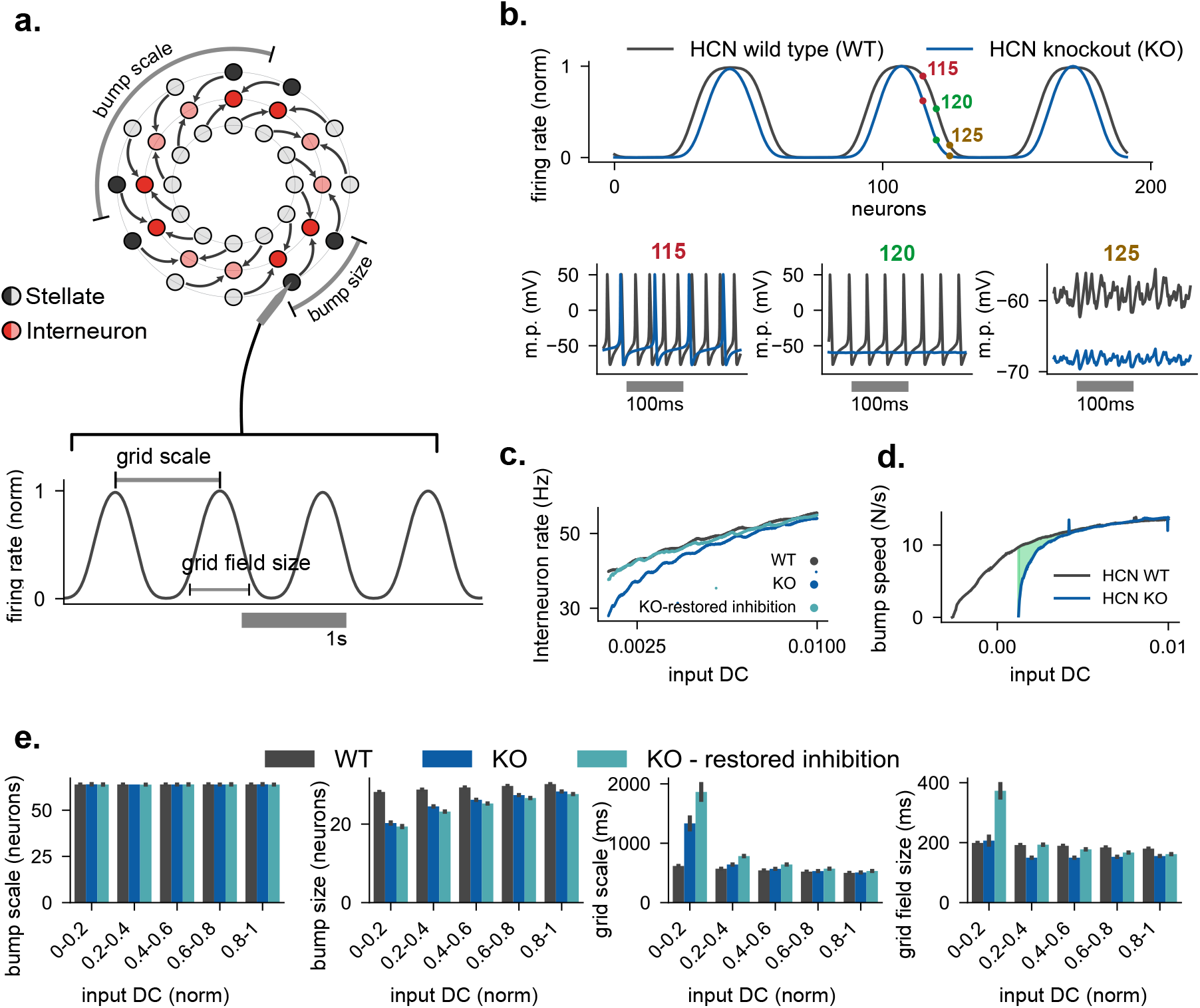
HCN reduces the velocity gain of the ring attractor. **(a) Grid scale vs bump scale.** Top, Bump scales and bump sizes are measured along the neurons. The bump scale is a measure of the distance between the centers of two consecutive bumps, and bump size is the number of neurons that constitute a single bump. Bottom, Grid scale and Grid field sizes are measured from recordings of a single neuron along the time (constant speed input) or distance axis. **(b) HCN KO shrinks bump sizes**. Top, Reduction in bump size in HCN knockout (blue). Bottom, Membrane potential of stellate cells at different locations on the bumps marked in the top panel. Note the drop in resting membrane potential in HCN knockout. At the edges of the bump, the knockout stellate cells are less excitable which leads to a reduction in the bump size. **(c) HCN KO lowers inhibition**. Average interneuron firing rates as a function of Input DC in HCN wild type (grey), HCN Knockout (blue) and HCN knockout after inhibition is restored (cyan). **(d) HCN modulates velocity gain**. Attractor speed as a function of input DC. In HCN knockout (blue) higher input DC is required to initiate bump movement, and for most of the input range the same input causes slower bump movement when compared to the wild-type. **(e)** Grid scale and Grid Field size as a function of Input DC (speed) in HCN wildtype (grey), HCN knockout (blue) and HCN knockout after inhibition is restored (cyan). The range of Input DC used for these simulations is highlighted in **d**. Bump scales are not affected by HCN knockout. However, Bump sizes shrink due to a drop in the resting membrane potential of stellate cells. Grid scales expand, and grid field sizes expand if the inhibition within the network is restored. In all four cases, the values converge at higher DC inputs.

HCN channels affect the excitability of neurons by raising the resting membrane potential and lowering the input resistance [Mishra and Narayanan, 2023]. HCN knockout also eliminates the depolarizing sag potential that is a characteristic property of stellate cells [Giocomo and Hasselmo, 2009]. The neurons are therefore slower to repolarize, causing a reduction in firing rates. Since interneurons receive excitatory input from stellate cells, a decrease in input from stellate cells would also decrease the firing rate of interneurons. However, interneuron activity is maintained after HCN knockout [Giocomo et al., 2011b]. To match the pre- and post-HCN knockout firing rates of interneurons as observed in experiments, we added a compensatory depolarizing input to the interneurons in our model (Figure 4c). Predictably, post-HCN knockout, we see a pronounced decrease in resting membrane potential, in turn, a reduction in the overall excitability and spiking activity of stellate cells. Reduced excitability meant that stellate cells near the border of activity bumps that receive higher inhibition relative to neurons at the center of the bump do not spike. This results in a reduction in bump size without a concomitant change in the bump scale, measured as the distance between neighboring bump peaks (Figure 4b,e). The reduced firing rates of the stellate cells that provide the directional offsets in the attractor cause a slowdown in the movement of the bump along the ring (Figure 4d). This reduction in attractor speed causes both grid scale and grid field expansion in our model (Figure 4e).

### 2.5 Cellular and network-based mechanisms for predictive and retrospective coding

The trajectory of an animal crosses the spatially circumscribed firing fields of multiple grid cells. Therefore, each trajectory can be mapped to a sequence of spikes by grid cells with overlapping grid fields. These sequences may exhibit either a predictive bias, representing sections of the trajectory ahead of the animal, or a retrospective bias, representing sections behind the animal [Ouchi and Fujisawa, 2024; Chaudhuri-Vayalambrone et al., 2023; De Almeida et al., 2012]. The magnitude and direction of these biases vary between layer II and layer III grid cells. Layer III grid cells display a strong predictive bias, extending up to 25% of the grid scales, while layer II grid cells exhibit a smaller predictive bias (~5% of the grid scale). Additionally, some layer II grid cells show a retrospective bias. Based on these observations, we propose two mechanisms that generate predictive and retrospective positional biases in our model network. Furthermore, we find that these mechanisms, one based on intrinsic cell properties and the other on network asymmetries, can explain the different bias magnitudes observed in layer II and layer III grid cells.

To understand the mechanism underlying the predictive bias, we examined the location of grid fields with and without HCN conductance. We compared the grid field locations when the animal traveled clockwise (rightward) to when it moved counterclockwise (leftward) (Figure 5a). We found a predictive bias in the direction of travel that extended up to~ 8% of the size of the grid field. When HCN channels were absent, we found that the direction of bias reversed to a retrospective bias, extending to 4% of the grid field. How does HCN give rise to a predictive bias? The HCN channel depolarizes the cell and is activated by inhibitory input from interneurons in the model network. Since HCN increases the cell’s excitability, it speeds up the onset of spiking, translating grid fields that are farther away to locations closer to the animal [Mishra and Narayanan, 2023].

**Figure 5.**
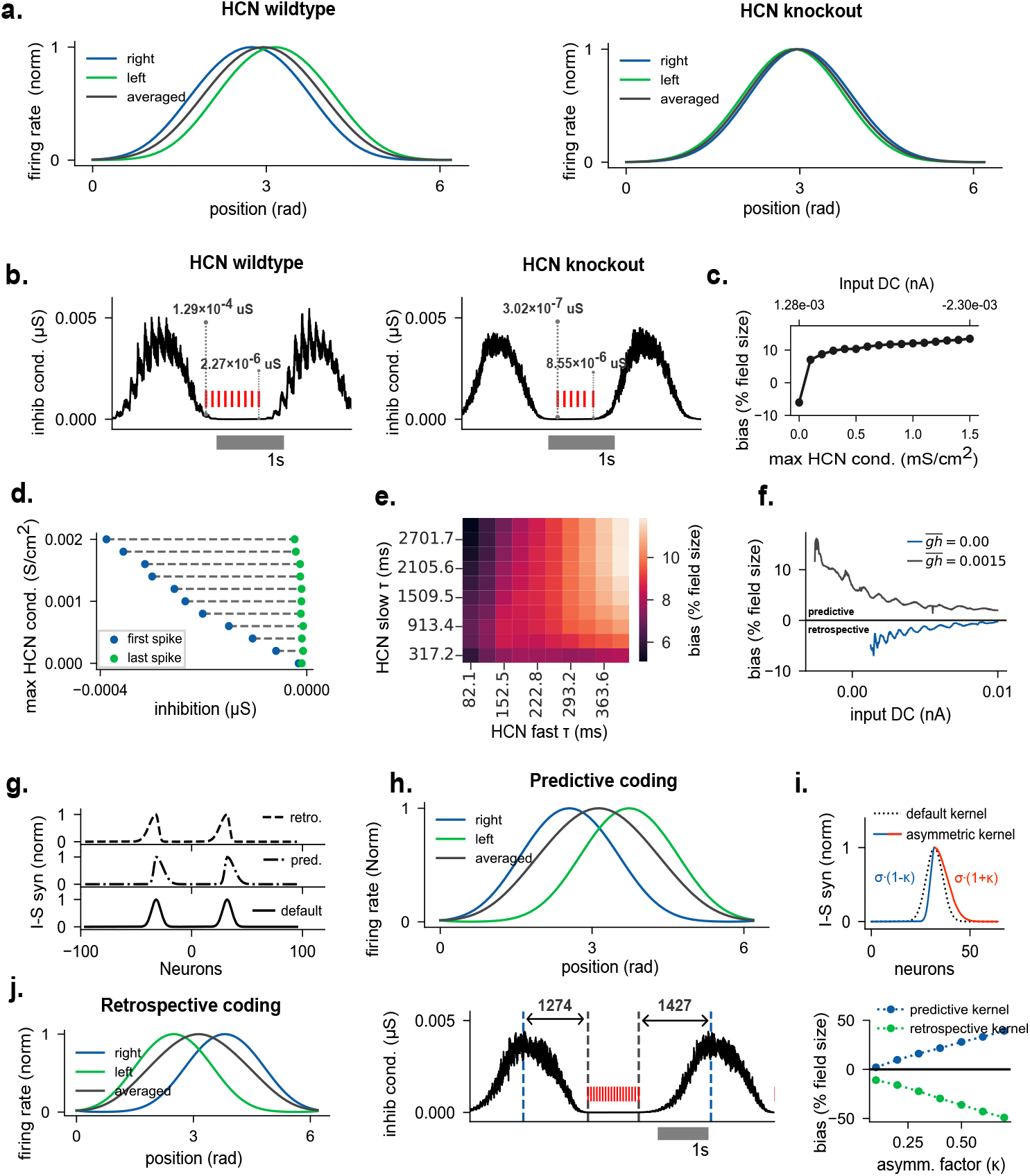
Predictive coding in the grid cell attractor model. **Neuron intrinsic predictive and retrospective coding** - **(a)** (left) Grid fields from a single stellate cell (averaged across consecutive fields) when the animal is moving either rightward (blue) or leftward (green). The gray curve is the right and left averaged grid field. (right) The same experiment in HCN knockouts abolishes predictive coding. **(b)** Comparison of inhibitory synaptic conductance received by a stellate cell in the presence (left) and absence (right) of HCN after a single pass through a grid field (in red). The annotations denote the values of the synaptic conductance at the start and the end of the field. **(c) HCN density.**Positional bias as a result of increasing HCN density. For each HCN density input DC was adjusted to maintain a constant bump velocity (*~* 30*N/s*) across all HCN density levels. **(d) Onset of first and last spikes**. The inhibitory synaptic conductance at the first and last spikes of grid fields is shown at increasing levels of maximal HCN conductance. **(e) HCN time scales**. The effect of both slow and fast time scales of the HCN conductance on the predictive bias. **(f) Speed dependence**. Positional bias measured as a function of increasing DC inputs (speed of the animal) in the presence and absence of HCN conductance. Positive values indicate predictive coding and negative values indicate retrospective coding. **Network based predictive and retrospective coding** - **(g)** The synaptic profiles of interneuron to stellate connections (clockwise ring) that are used to generate predictive and retrospective bias in the absence of HCN conductance. A mirrored version of the same connection (not shown) is projected to the counterclockwise ring. **(h)** Top, Network-based predictive bias. As in a, grid fields are shown as the animal is moving in left (green) or right (blue) directions. The gray curve is the right and left averaged grid field. Bottom, Inhibitory synaptic conductance that is received by the same stellate cell. Note the duration of the inhibition is longer at the end of the field. **(i)** Top, One lobe of the synaptic profile in g (middle) is shown. Asymmetry is generated by adjusting the asymmetry factor *κ*, which extends the Gaussian on the right side of the peak and shrinks it on the left side. Bottom, The magnitude of the bias as a function of the asymmetry factor in the predictive and the retrospective network. **(j)** Retrospective bias generated by using the connectivity depicted in the top panel in F.

As we increased the conductance of the HCN channel, the cell became more excitable and could spike at higher levels of inhibition. The overall inhibitory drive at which the stellate cell spiked was nearly 10 times higher for the HCN conductance values seen in wild-type [Acker et al., 2003; Dickson et al., 2000], compared to when the HCN channel was absent (Figure 5d. See annotations in Figure 5b). Therefore, the onset of the first spike occurred earlier, leading to a larger predictive bias at higher HCN conductance values (Figure 5d).

Grid cells are arranged in modules along the dorsoventral axis of the MEC with increasing grid scales and grid field sizes [Stensola et al., 2012]. Changes in grid scale are accompanied by concomitant changes in the time scale of HCN-dependent depolarizing currents that increase along the dorsoventral axis [Giocomo and Hasselmo, 2008]. We tested whether the time scales of HCN can affect the predictive bias observed in our model. We modeled the HCN conductance with a slow and fast time scale [Acker et al., 2003]. Figure 5e shows a consistent increase in predictive bias as both the slow and fast time scales of HCN are increased. Since the length scale of grid fields co-vary with HCN time scales along the dorsoventral axis, the observed positive correlation between grid scales and the magnitude of predictive bias is likely a result of increasing HCN time scales as seen in our model [Chaudhuri-Vayalambrone et al., 2023; Ouchi and Fujisawa, 2024]. The range of shift distances reported across grid modules [Chaudhuri-Vayalambrone et al., 2023; Ouchi and Fujisawa, 2024] is consistent with a mechanism in which larger, more ventral grid modules, which are associated with longer HCN time constants, produce proportionally larger predictive shifts, as our model predicts; a direct quantitative comparison will require matching the HCN time-constant range explored in the model to the specific grid modules recorded experimentally.

The magnitude of the observed bias (predictive or retrospective) is speed dependent, with the bias being highest at low forward speeds of the rat and minimal at high speeds [Chaudhuri-Vayalambrone et al., 2023]. We found that this was also the case in our model network for different HCN conductance values for a range of speeds (Figure 5f). The extent of predictive and retrospective bias seen in our model by varying the time scales and conductances of the HCN channel within an experimentally observed range is consistent with that seen in layer II stellate cells.

Layer III grid cells, in contrast, show a predictive bias that extends over ~25% of a grid field. Changes in the conductance and time scales of the HCN channel are not sufficient to generate these observed biases.

To reproduce the larger predictive bias observed in the layer III grid cells, we introduced asymmetries in the interneuron to stellate connectivity (Figure 5g). Consider the inhibitory input from the interneuron ring to the outer ring of stellate cells (Figure 1d) with grid fields that move clockwise. In our simulations thus far, inhibitory input to the stellate cells followed a symmetric bimodal Gaussian function (Figure 5g, bottom panel). To amplify the predictive bias, we introduced an asymmetry around each peak of the bimodal Gaussian function (Figure 5g middle panel, Figure 5i).

We implemented a mirrored version of the same connectivity (not shown in Figure 5) that inhibited the inner ring of stellate cells with grid fields that moved in the counterclockwise direction. As the animal traversed the ring clockwise, it periodically inhibited presynaptic interneurons (Figure 5h bottom panel). The asymmetric connectivity of the inhibitory network meant that inhibitory input to each stellate cell decayed rapidly, and the external constant depolarizing drive to stellate cells was able to elicit a spike at earlier times than the symmetric network. Note that in these simulations, we did not include an HCN conductance in the stellate cells so that one could disambiguate the effects of predictive spiking due to rebound spikes from those due to network asymmetries. We were able to change the extent of predictive bias by changing the degree of asymmetry in inhibitory connectivity (Figure 5i). In addition, we were also able to generate a retrospective bias in the model network by switching the direction of the asymmetry around the peak of each Gaussian function (Figure 5j).

## 3 Discussion

We developed a network model for grid cells using conductance-based neurons and incorporating realistic membrane currents that are seen in stellate cells and inhibitory interneurons of the medial entorhinal cortex. Our model can integrate complex trajectories accurately over behavioral timescales. To achieve accurate path integration a linear relationship between the speed of the animal and the movement of attractor is necessary. We achieved this by introducing an invertible mapping between speed and the amplitude of the depolarizing input used to drive the stellate cells. Consistent with experiments, we also show that grid fields arise due to disinhibition of stellate cells that results in a ramp-like increase of the membrane potential as the rat enters a grid field [Domnisoru et al., 2013]. As a consequence of this disinhibition, HCN conductance is maximum before the onset of the grid fields. Higher HCN conductance increases the excitability of stellate cells and hence advances the onset of the fields. A key finding of this study is how this can give rise to an “intrinsically generated” predictive bias in the position representation of stellate cells. Alternatively, a larger network-based predictive bias can also be generated by introducing asymmetries in the interneuron to stellate connectivity. In addition to introducing a positional bias, HCN can also adjust the velocity gain of the attractor network, leading to changes in grid scaling and field sizes that have previously been observed in experiments [Giocomo et al., 2011b].

Predictive grid cells recorded *in vivo* are phase-locked to the local theta rhythm [Ouchi and Fujisawa, 2024]. Our model introduces theta oscillations as a drive to the MEC network (Figure 1a), and we find that, for sufficiently large theta-modulated input amplitudes, stellate cell spiking becomes entrained to the theta rhythm, consistent with this phase-locking. A full characterization of the relationship between this entrainment and the predictive and retrospective biases described above is beyond the scope of the present study and is left for future work.

Most attractor models of grid cells achieve accurate path integration by linearly modulating the network’s input, resulting in a proportional shift of the attractor along a specific direction. The activity of a grid cell at any given location depends only on the animal’s current position and is unaffected by its past trajectory. Consequently, the differential equations that govern the activity of the network must be linear functions of velocity, aligned with the way that attractor models incorporate the animal’s movement. However, the firing rate of stellate cells changes nonlinearly in response to input [Fransén et al., 2004]. The movement of the attractor bump is influenced by various intrinsic and synaptic factors, causing it to respond nonlinearly to an external drive. Furthermore, the response of speed cells to velocity can range from simple linear relationships to more complex nonlinear dependencies[Kropff et al., 2015; Hinman et al., 2016]. Even speed cells with a linear response to speed exhibit diverse slopes and intercepts. Given the diversity of speed cell responses, we predict that the velocity input to grid cells must be modulated in a manner that linearizes the eventual output of the grid cell network. The presence of a modulatory mechanism allows the spatial representation to be flexible without changing the structure of the grid cell network. For example, the hexagonal pattern of grid cell responses can be flexible when measured in different environments geometries [Krupic et al., 2015], or with different degrees of familiarity within a given environment [Barry et al., 2012]. If grid cell activity distortions were a result of long-term synaptic plasticity in the attractor network, it would prevent the formation of a stable metric, an allocentric coordinate system across different environments. However, if the environmental geometry and the task structure influence which speed cells serve as input to grid cells or modulate the speed input to grid cells; the presence of alternate tuning curves would allow temporary distortions to grid cell responses without effecting long-term changes to the structure of the attractor network. Speed cells themselves can carry a temporally offset signal: a subset of speed-tuned cells in the MEC represent the animal’s running speed with a predictive or retrospective offset relative to the true instantaneous speed [Kropff et al., 2015]. Our model represents the population speed signal as a single, instantaneous depolarizing drive I = f(s) and does not currently capture this heterogeneity. If even a fraction of the depolarizing drive to stellate cells were itself temporally offset in this way, it could shift the magnitude, or potentially the sign, of the retrospective bias that we attribute here to network asymmetry, providing an additional, speed-coding-dependent route to retrospective coding that could act alongside the intrinsic and network mechanisms we describe. Disentangling the relative contributions of speed-signal timing and network asymmetry to retrospective coding is a promising direction for future modeling work.

HCN channels in MEC stellate cells show a rich dynamical repertoire with systematic changes in the dynamics of the subthreshold membrane potential and the spiking properties of the cell in addition to synaptic integration times [Nolan et al., 2007; Garden et al., 2008] along the dorsoventral axis of the MEC. Although *in vivo* recordings show that HCN modulates the grid scale and grid field sizes, it is unclear how HCN contributes to spatial computation in grid cells [Giocomo et al., 2011b]. Here, we show that the HCN channel biases grid fields along a direction opposite to the movement of the animal. As a consequence, we predict that knocking out HCN channels in layer II stellate cells will eliminate the HCN-dependent predictive bias while retaining a smaller retrospective bias, which arises from the directed asymmetric excitatory input from stellate cells to inhibitory interneurons. This asymmetric connectivity determines the direction of propagation of the attractor bumps. In contrast, we model input from interneurons to stellate cells using a symmetric connectivity kernel. Asymmetries in these connections introduce a prospective (or retrospective) bias without affecting the direction of propagation of the attractor bump. The network asymmetry-induced bias does not depend on HCN channels and can be tuned to vary across a broad range. This is consistent with the observation that non-stellate cells of the entorhinal cortex lack the HCN dependent electrophysiological properties that give rise to post-inhibitory rebound spikes [Alonso and Klink, 1993; Schmidt-Hieber and Häusser, 2013]. In layer II, the mean predictive bias is*~* 5% of the grid scale. In contrast, the mean predictive bias in layer III is nearly 25% of the grid scale, indicative of a network asymmetry-induced origin. The relative magnitude of the predictive biases in layers II and III raises an interesting question - Does the asymmetry of inhibitory input to layer III grid cells originate in layer II interneurons, or is it due to interneurons within layer III? The latter could be evidence of a separate attractor network in layer III of mEC. However, optogenetic stimulation of layer II pyramidal cells causes di-synaptic inhibition in layer III pyramidal cells, suggesting that the predictive grid cells in layer III MEC are a direction-biased readout of the attractor network in layer II[Zutshi et al., 2018].

The present model treats all stellate cells as belonging to a single grid module with a fixed spacing; it does not include a mechanism for generating the discrete, quantized modules observed along the dorsoventral axis [Stensola et al., 2012]. Different grid scales can nonetheless be reproduced in our framework by changing the spatial extent of lateral inhibition, and the same HCN-dependent expansion of grid scale and field size described above would be expected to apply within each such module. Extending the model with a graded, dorsoventral variation in lateral inhibition, along the lines proposed for module self-organization [Kang and Balasubramanian, 2019; Khona et al., 2025], is a natural direction for future work.

As in most continuous attractor network models of grid cells, several elements of our network’s connectivity, including the offset excitatory projections from stellate cells to interneurons that set the direction of attractor propagation, and the asymmetric interneuron-to-stellate connectivity that generates the layer III-like predictive bias, are constraints that are not derived from a fully specified, data-driven wiring diagram of the MEC. There are marked differences between the predictive coding of layer II and layer III grid cells, with the predictive bias in layer III roughly five-fold larger than in layer II. Our model predicts that these differences arise from an asymmetric directional skew in the synaptic weights between interneurons and stellate cells, present in layer III and absent, or much weaker, in layer II. Such a difference in synaptic weight or connection probability should become evident in paired recordings or connectomic reconstruction of interneuron-stellate synapses. This asymmetry has not yet been demonstrated directly, though recent anatomical work showing that stellate cell-interneuron synapses are organized at the spatial scale of individual grid-cell clusters [Huang et al., 2024] is at least consistent with the kind of spatially patterned connectivity our model requires (schematic figure 8 in Huang et al. [2024]).

In summary, our work demonstrates that a continuous attractor network built from biophysically detailed, conductance-based stellate cells and inhibitory interneurons can accurately path integrate while reproducing a range of phenomena observed in the MEC, including the depolarizing membrane potential ramp within grid fields, HCN-dependent changes in grid scale, and the predictive and retrospective biases of grid cell firing. Central to these results is the finding that predictive coding in the MEC need not arise from a single source. Instead, it can emerge from two dissociable mechanisms: an intrinsic, cell-autonomous mechanism, in which the rebound dynamics of the HCN conductance advance the onset of spiking and generate the modest predictive bias of layer II stellate cells, and a circuit-level mechanism, in which asymmetries in inhibitory connectivity produce the larger predictive bias characteristic of layer III. That the intrinsic mechanism scales with the HCN time constants known to vary along the dorsoventral axis offers a concrete, experimentally testable link between the biophysics of a single channel and the spatial reach of the neural code. By grounding predictive coding in the wetware of the MEC, our model bridges single-neuron biophysics and population-level spatial computation.

## 4 Methods and Materials

### 4.1 Neuron and Synaptic Models

Stellate cells of the medial entorhinal cortex (MEC) were modeled as in [Acker et al., 2003], a reduced version of the more detailed conductance-based stellate cell model of [Fransén et al., 2004]. The conductance-based model shows postinhibitory rebound, subthreshold oscillations, depolarizing sag, and membrane resonance. As shown in Equation 7, cells are endowed with transient sodium, potassium, and leak currents seen in the standard Hodgkin-Huxley model. Additionally, the model includes hyperpolarization activated cation currents (Ih) and persistent sodium currents (Nap), which are active in the subthreshold regime. These currents are responsible for the properties of the stellate cells mentioned above.

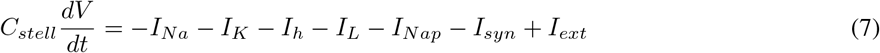

Fast-spiking inhibitory interneurons from [Wang and Buzsáki, 1996] were utilized to model feedback inhibition of the continuous attractor network (Equation 8). These cells were also theta modulated through a conductance-based oscillatory current to model theta oscillatory input from the medial septum to inhibitory interneurons of the MEC.

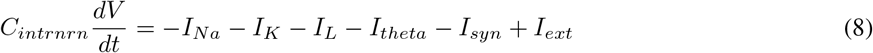

Both excitatory and inhibitory synapses were modelled using NEURON’s built-in event-based Exp2Syn. Exp2Syn can model both AMPA and GABA_*a*_ type synapses as a difference of two exponential with different time constants for rise and decay. All models were simulated in parallel in the NEURON simulation environment using its python interface [Carnevale and Hines, 2006].

#### 4.1.1 2D model

The topology of the 2-D model was similar to that of Kang and DeWeese [2019] with the primary differences being the presence of mutual inhibition within the interneuron sheet, no mutual excitation among the stellate sheet, and each sheet is connected along the edges to form a torus. For each sheet, neurons were placed on a 2D cartesian plane along integer locations. To achieve toroidal connectivity, the distance between two neurons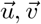 is given by:

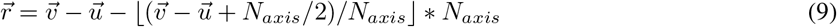

where, N_*axis*_ is number of neurons along one axis of the sheet and x is the floor operation. The size of the network, five sheets of 71× 71 neurons for the 2-D model, and three rings of 192 neurons for the reduced 1-D model, was chosen to balance biophysical realism against the computational cost of simulating conductance-based neurons over the many-seconds-to-minutes-long trajectories, and the extensive multi-parameter sweeps, required to characterize the network. Periodic boundary conditions and radially symmetric connectivity were used throughout to minimize edge and finite-size effects on the resulting attractor dynamics.

### 4.2 Path Integration in 1D

Position estimation in a population of grid cells within a module with grid scale λ, is equivalent to determining where the rat is in between two grid fields of the same cell. In other words, which grid phase is active at the current position [Fiete et al., 2008]. To assign a phase to each stellate cell, we uniformly distribute all stellate cells between two successive bumps along a length λ (of arbitrary spatial units). Then, for n neurons between two bumps, the phase of each neuron j is given by 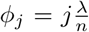. To decode positions from the activity of stellate cells we utilized a population vector-based decoder from [Stemmler et al., 2015]. The position x(t) is given by,

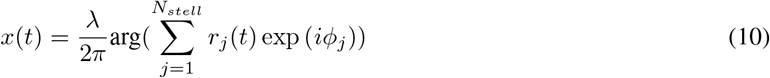

Where, r_*j*_(t) is the instantaneous firing rate of neuron j and arg gives the angle of the population vector in the complex plane. We assumed that the rat is moving on a circular track with a radius of 1. Therefore, in all our simulations, λ was set to 2π.

### 4.3 Membrane potential ramp and theta amplitude envelope

The ramp and theta amplitude envelopes were measured based on [Domnisoru et al., 2013]. The spikes were removed from the membrane potential, and the edges were connected using linear interpolation to obtain a spike-free potential. The resulting signal was passed through a low-pass filter with a cut-off at 3 Hz, and the mean spike-free potential was subtracted from it to produce the ramps. The mean was calculated for only out-of-field values of the spike-free potential. For the theta-amplitude envelope, the spike-free potential was band-pass filtered from 4-12 Hz. The magnitude of the analytical signal computed using the Hilbert transform of the filtered signal gave the theta amplitude envelope.

### 4.4 Predictive and retrospective coding

To calculate the positional bias, we simulated the forward and reverse movement of the animal for 40 seconds each. The decoded positions were estimated after 3 seconds in each direction to allow the network to stabilize. Since we were only concerned with the relative difference in the positions between the right (clockwise) and left (counterclockwise) directions, a reference “current position” was calculated using a low-pass-filtered activity from the interneuron ring. This filtering was done to avoid any positional bias introduced by the stellate cell rings. The interneuron ring’s firing rate does show a small asymmetry opposite to the direction of travel; however, unlike the stellate cell rings, this asymmetry is not accompanied by a translation of the interneuron population’s positional tuning, and we therefore used the low-pass-filtered interneuron activity as an unbiased estimate of the animal’s instantaneous position. The timing of stellate spikes was then mapped to these positions to give the firing field of the stellate cells as a function of position. Figure 5a, h and j were then produced after smoothing and averaging the estimated firing fields across consecutive fields in individual movement directions. For Figure 5e simulations were run for 360 s. HCN time constants were calculated by incrementing k_*f*_, k_*s*_ in,

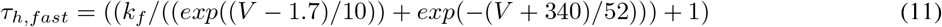

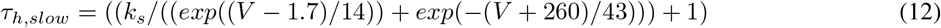

The range for k_*f*_ and k_*s*_ was chosen to match the HCN time constants measured in [Giocomo and Hasselmo, 2008]. The axes labels in Figure 5e denote the peak time constant for each value of k.

### 4.5 Simulation environment and code availability

The network was simulated using the NEURON simulator (version 8.2) [Carnevale and Hines, 2006] with its Python API, employing a parallelized implementation. Smaller-scale exploratory simulations were run on a 40-core work-station with 512 GB memory, while larger-scale grid searches were distributed on a High-performance cluster utilising up to 12 nodes. Code for simulations with corresponding notebooks and documentation is available at https://github.com/assisilab/GridCellsCond

## 6 Acknowledgments

We thank members of Assisi and Nadkarni labs at IISER Pune for useful discussions. We also acknowledge the National Supercomputing Mission (NSM) for providing the computing resources of ‘PARAM Brahma’ at IISER Pune, which is implemented by C-DAC and supported by the Ministry of Electronics and Information Technology (MeitY) and Department of Science and Technology (DST), Government of India

